# Generation and mutational analysis of a transgenic mouse model of human SRY

**DOI:** 10.1101/2021.03.04.433906

**Authors:** Ella Thomson, Liang Zhao, Yen-Shan Chen, Enya Longmuss, Ee Ting Ng, Rajini Sreenivasan, Brittany Croft, Xin Song, Andrew Sinclair, Michael Weiss, Emanuele Pelosi, Peter Koopman

**Affiliations:** Institute for Molecular Bioscience, The University of Queensland, Brisbane, QLD 4072, Australia; Centre for Clinical Research, The University of Queensland, Brisbane, QLD 4072 Australia; Department of Biochemistry & Molecular Biology, Indiana University School of Medicine, Indianapolis, IN 46202, USA; Murdoch Children’s Research Institute and Department of Paediatrics, University of Melbourne, Melbourne, VIC 3052, Australia

**Keywords:** Sry, sex determination, DSD, structure-function, mouse models, CRISPR

## Abstract

SRY is the Y-chromosomal gene that determines male sex development in humans and most other mammals. After three decades of study, we still lack a detailed understanding of which domains of the SRY protein are required to engage pathway of gene activity leading to testis development. Some insight has been gained from the study of genetic variations underlying differences/disorders of sex determination (DSD), but the lack of a system of experimentally generating SRY mutations and studying their consequences in vivo has limited progress in the field. To address this issue, we generated a mouse model carrying a human *SRY* transgene able to drive male sex determination in XX mice. Using CRISPR-Cas9 gene editing, we generated novel genetic modifications in each of *SRY*’s three domains (N-terminal, HMG box, and C-terminal) and performed detailed analysis of their molecular and cellular effects on embryonic testis development. Our results provide new functional insights unique to human *SRY* and the causes of DSD, and present a versatile and powerful system in which to demonstrate causality of *SRY* variations in DSD, to functionally study the *SRY* variation database, and to characterize new pathogenic *SRY* variations found in DSD.

## Introduction

Development of the testis in males is controlled in most eutherian mammals, including humans, by the Y-chromosome gene *SRY* (Koopman et al., 1991). Expressed within the somatic supporting cells of the bipotential gonad, the SRY protein activates expression of *SOX9* through binding to enhancers including TESCO, eSR-A/Enh13 and eALDI (Albrecht and Eicher, 2001, Gonen et al., 2017, Croft et al., 2018). SOX9 subsequently upregulates the testis-specific developmental pathway while suppressing pro-ovarian gene expression. In the absence of *SRY*, *SOX9* is not activated within these cells, and reciprocal ovarian-specific pathways supervene.

Despite its critical role in human evolution and biology, our understanding of the biochemical roles of SRY, and of what parts of the protein are required to carry out those roles, remains limited. While these issues have been addressed in some detail in mice (Zhao et al., 2014a), the low level of sequence conservation between human and mouse SRY, and the markedly different topographies of the two proteins (Figure 1A) renders studies in mice of limited value in understanding the structure-function relationship in human SRY.

**Figure 1:**
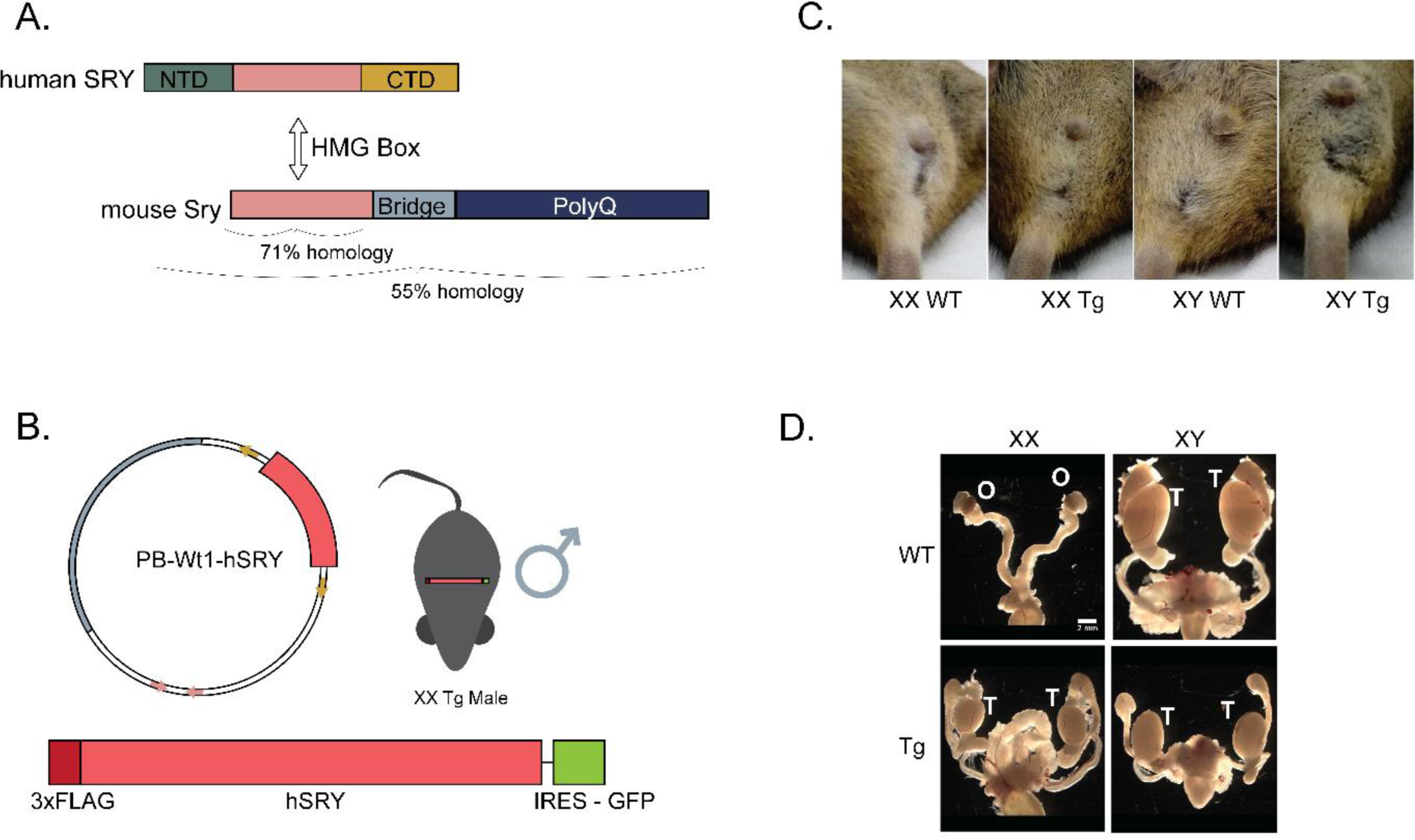
Generation of the human *SRY* sex reversed mouse line. **A.** The *Sry* gene in human and mouse is highly divergent. Both contain the HMG box, encoding the DNA binding domain (pink) which is 71% homologous between the species. Human *SRY* has an N-terminal (green) and C-terminal domain (yellow) which are not seen in the mouse. **B.** The PB-Wt1-hSRY construct was generated with a 3xFLAG-hSRY gene, followed by a GFP reporter. This was inserted into the mouse genome, resulting in XX Tg mice appearing as male. XY Tg mice are used to breed for propagation of the line. **C.** The external reproductive organs of WT and Tg 6-week-old WT and Tg mice. XX WT mice show the urethral and vaginal orifice, while XX Tg and XY Tg show a penis and scrotal sac consistent with XY WT anatomy. **D.** The internal reproductive organs of WT and Tg 6-week-old XY and XX mice. WT XX mice develop ovaries, while XX Tg and XY Tg mice show testis development. Both XX and XY Tg have smaller testis than the XY WT control. T, testis; O, ovary. Scale bar, 2mm.

Much of what is known about human SRY function, and the corresponding functional domains, has come from study of pathogenic variants found in human differences/disorders of sex development (DSD). Variants of SRY are the most common cause of 46,XY DSD, usually manifested as partial (PGD) or complete gonadal dysgenesis (CGD) (McCann- Crosby et al., 2014). The majority of *SRY* variants associated with 46,XY DSD are found within the HMG box, which encodes the DNA-binding domain (Supplementary Table 1). Very few pathogenic variants have been described within the N-terminal domain (NTD), including truncations leading to a loss of the HMG box (Brown et al., 1998, Veitia et al., 1997, Scherer et al., 1998, Xue et al., 2019); such variants are uninformative regarding whether the N-terminal domain is required for any specific SRY functions, or what those functions might be. Similarly, only six DSD-causing variants have been identified in the C-terminal domain (CTD) (Shahid et al., 2004, Sánchez-Moreno et al., 2009, Baldazzi et al., 2003, Shahid et al., 2005, Tajima et al., 1994, Özen et al., 2017). Because the CTD has been implicated in transcriptional trans-activation (Dubin and Ostrer, 1994), it is assumed that these variants interfere with this function of the SRY protein. It remains unknown whether the NTD and/or CTD play any roles in protein stabilisation, partner protein binding, target specificity or other biochemical mechanisms.

Due to the lack of an appropriate *in vivo* system to test clinical impact of variants in *SRY*, at present *in vitro* and cell-based experiments have provided the only way to assess SRY activity. However intriguing, such experiments do not directly test the capacity of an SRY variant to activate the male developmental program in the mammalian bipotential gonad. To circumvent this limitation, we developed a transgenic mouse model harbouring the human *SRY* gene (*hSRY*), which can then be interrogated *in vivo* by creating predetermined modifications using CRISPR. In this system, the transgene triggers formation of testes in XX mice, with gene-expression patterns reflecting those of typical XY testes. We showed that specific modifications to the N- and C-terminal and HMG domains of the *hSRY* transgene resulted in complete or partial loss of SRY function. We demonstrate the utility of this strategy as a genetic tool for determining SRY structure-function relationships pertinent to human sex development and developmental difference.

## Results

### Generation of a transgenic human SRY mouse model

We utilised the Gateway enhanced PiggyBac system (Zhao et al., 2014b) to generate a novel transgenic mouse model. The transgene was composed of an N-terminal 3x FLAG-tagged human *SRY* followed by an internal ribosome entry sequence (IRES) and the gene encoding enhanced green fluorescent protein (eGFP) (Figure 1B). It uses the regulatory regions of *Wt1* — expressed in the pre-Sertoli cells of the bipotential gonad (Pelletier et al., 1991, Armstrong et al., 1993, Chen et al., 2017) — to drive expression of the transgene specifically to the developing urogenital ridge. This approach has been used successfully to express and study mouse *Sry* structure and function (Zhao et al., 2014a).

Transgenic mice were generated by pronuclear microinjection of the PB-Wt1-hSRY construct (Figure 1B). Breeding of a transgenic (Tg) XY male founder with a wild type female confirmed germline transmission. Splinkerette PCR analysis showed integration of a single copy of the *hSRY* transgene in an intergenic region of chromosome 8 with no known functions and no highly conserved regions (Supplementary Figure 1).

### XX transgenic mice develop as males

Both XY and XX transgenic progeny appeared phenotypically male and indistinguishable from XY wild type littermates (Figure 1C). Examination of the internal reproductive organs revealed that XX Tg animals developed an entire male reproductive system, including testes and vasa deferentia rather than ovaries and uterus (Figure 1D).

Although XY Tg testes appeared smaller than wild type controls, histological analysis showed that the seminiferous tubules of XY Tg testes were smaller than their wild type counterparts and had fewer sperm (Figure 2A). However, this did not affect the fertility of XY Tg males, which were successfully used as breeders for the propagation of the mouse line.

XX Tg mice were sterile, as expected owing to the lack of Y-chromosome genes necessary for spermatogenesis, and consistent with the phenotype of human 46,XX DSD. Their gonads were smaller than wild type testes; the seminiferous tubules lacked germ cells and displayed the typical architecture of streak XY gonads (Figure 2A). Immunofluorescence (IF) for the germ cell marker MVH confirmed that, in contrast to XY wild type and XY Tg testes, and the follicles of XX wild type ovaries, XX Tg samples did not show any visible signal (Figure 2B).

**Figure 2:**
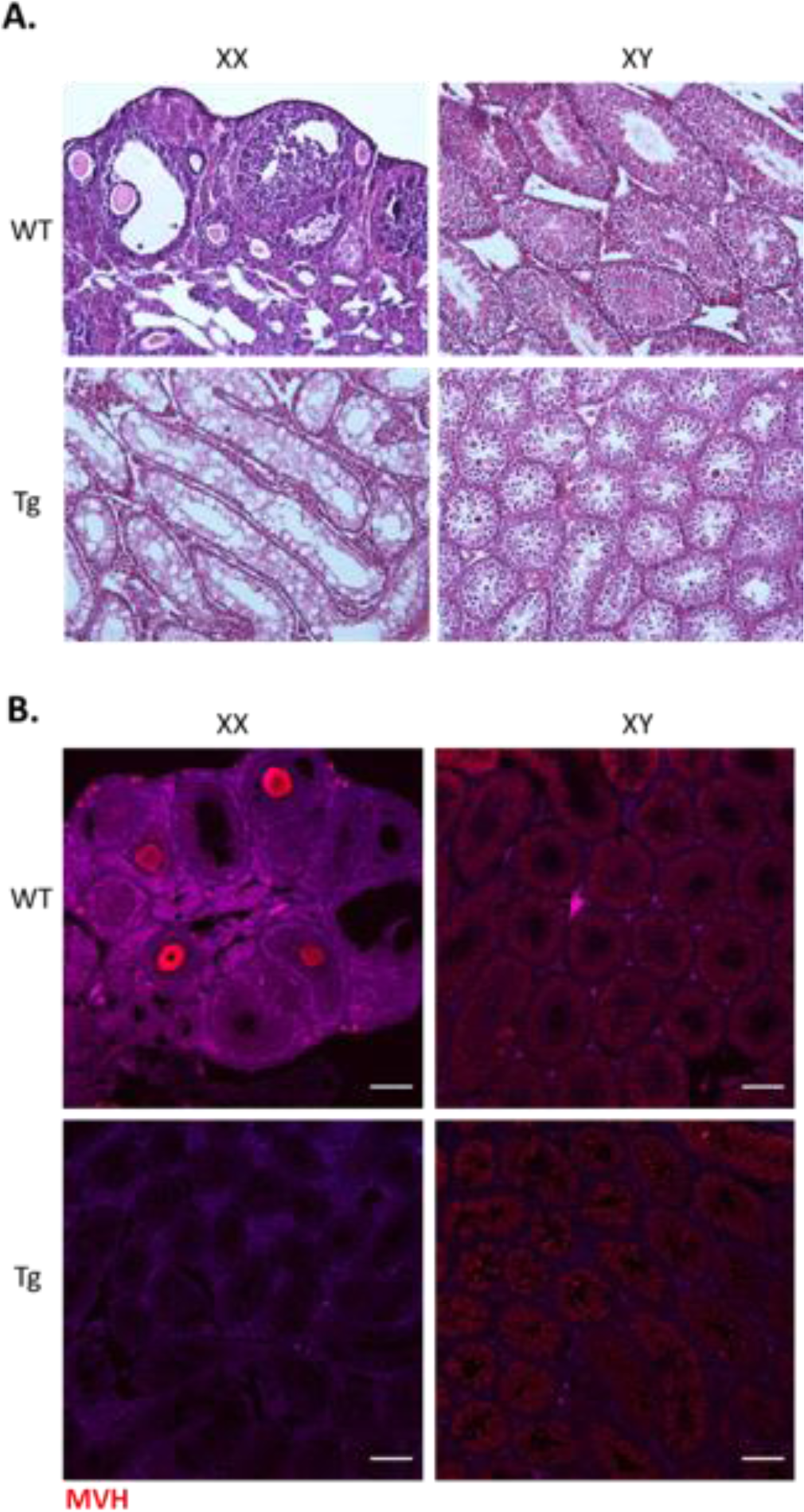
Postnatal analysis of the *hSRY* mouse line. **A.** Morphology of 6-week postnatal gonads was assessed via H & E staining. XX and XY samples showed typical ovarian and testicular morphology. XX Tg samples appear testicular however no sperm are visible, and dysgenesis of the testis cords is apparent. XY Tg samples show a typical male testicular morphology. **B.** MVH staining of the postnatal sections identifies germ cells within the tissue. XX WT samples show staining within the ovarian follicles, and XY WT and XY Tg samples show MVH expression within testis cords. No staining is evident within the XX Tg samples. Scale bar, 100 μm.

### Human SRY is expressed by the supporting cells of the developing testis

To evaluate expression of *hSRY* at 11.5dpc (when endogenous mouse *Sry* is expressed), expression of the FLAG protein was analysed by IF together with the somatic cell marker GATA4 and the germ cell marker MVH. FLAG expression was detected specifically within the somatic supporting cells of the gonad only in transgenic animals (Figure 3A). No expression of FLAG was seen within the primordial germ cells.

**Figure 3:**
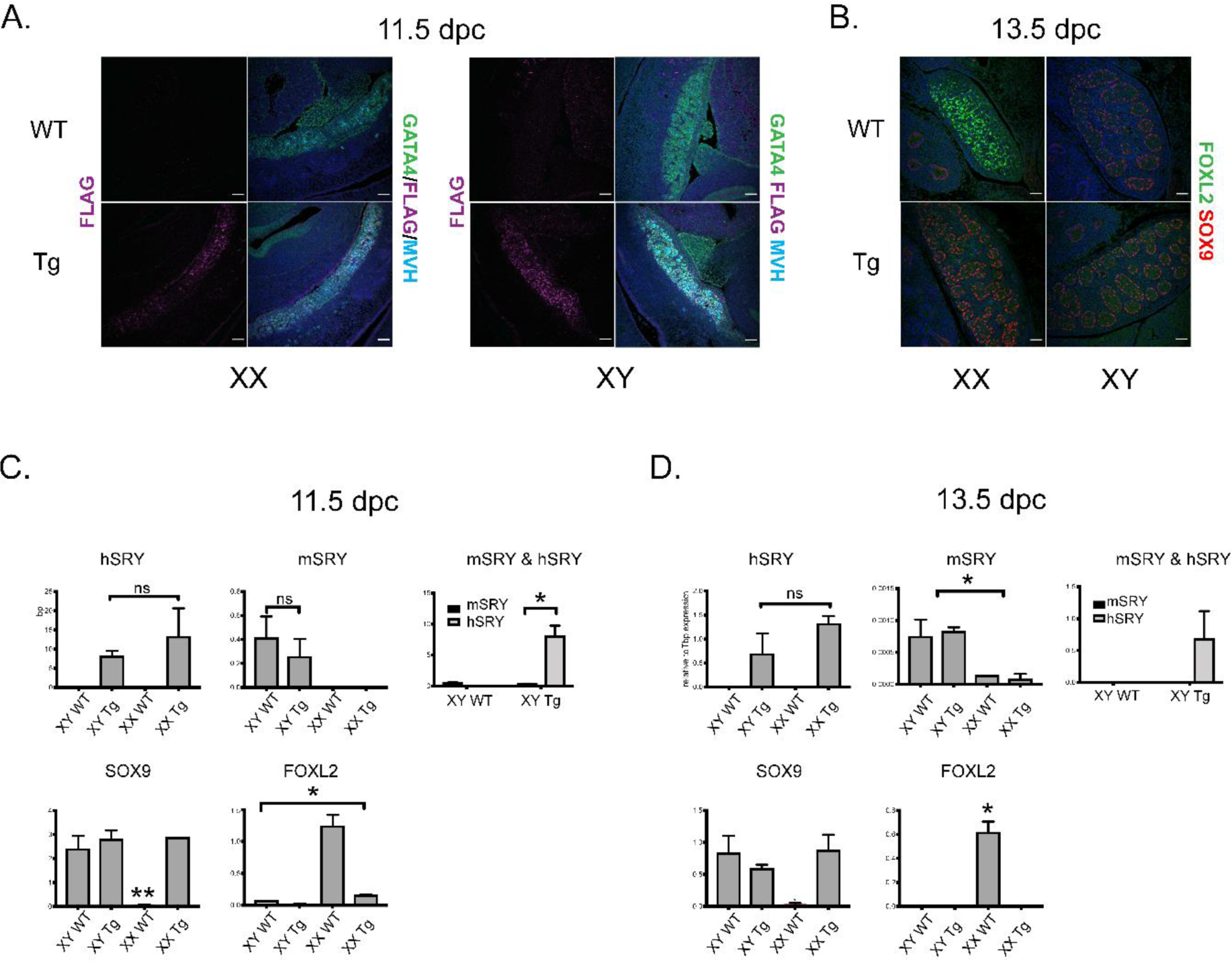
Gene Expression Analysis of *hSRY* mouse line. **A.** Expression of FLAG, GATA4 and MVH was assessed via IHC in 11.5 dpc gonads. Expression of FLAG can be seen only in Tg samples, with no expression visible in XX WT or XY WT samples. GATA4 marks the somatic cells of the gonad, and MVH marks the germ cells. Similar levels of each are visible throughout all genotypes. Bar = 50 μm. **B.** Sertoli cell marker (SOX9) and granulosa cell marker (FOXL2) were analysed in 13.5 dpc samples. XX WT samples show extensive FOXL2 staining with no SOX9 visible. Both XX Tg and XY Tg samples show similar staining patterns XY WT with SOX9 staining throughout testis cords. Bar = 50 μm. **C.** Expression levels of *hSRY, mSRY, Sox9* and *FoxL2* in were measured in XY WT, XY Tg, XX WT, XX Tg gonads at 11.5 dpc. **D.** Expression level of same markers at 13.5 dpc. In both 11.5dpc and 13.5dpc samples, *hSRY* is expressed within both XY and XX Tg lines, and this is expressed at levels much higher than the endogenous mouse *Sry* in XY lines. *Sox9* expression with XX and XY Tg lines at both timepoints is consistent with XY WT male expression. *Foxl2* expression in XX WT is significantly higher than all other genotypes, however XX Tg mice at 11.5 dpc show higher levels than XY samples.

### XX transgenic mice show male gene expression patterns in the gonad

When *Sry* is expressed in the bipotential gonad of a developing XY male, it upregulates *Sox9*, which in turn suppresses the ovary specific *Foxl2*. To investigate whether the *hSRY* transgene expression in fetal gonads induced gene expression similar to that of mouse *Sry* (*mSry*), we performed IF for SOX9 and FOXL2 at 13.5 dpc, after the gonads have differentiated (Figure 3B). We also performed qRT-PCR analyses of embryonic gonads at both 11.5 dpc (during the period of *mSRY* expression) and 13.5 dpc (Figures 3C, 3D). Both analyses showed that the expression pattern of *Sox9* and *Foxl2* was similar between XX and XY Tg gonads and wild type XY samples at fetal stages.

As the transgene was not driven by the endogenous *mSry* promoter, we determined the difference in expression between *mSry* and *hSRY*. qRT-PCR showed that *hSRY* was expressed at higher levels than *mSry* (Figures 3C, 3D). Further, *hSRY* was still expressed in the gonad at 13.5dpc, after *mSry* turns off. This extended expression likely has no functional consequence, because the ability of *Sry* to activate the testis developmental program is thought to depend only on its expression within a critical time window around 11.5 dpc.

### Proof of Principle: Deletion of the HMG box (“HMG-Del” mutation)

To understand the effects of genetic variants that could inactivate the human SRY, we used CRISPR to edit the transgene *in vivo*. CRISPR components were electroporated into single cell mouse zygotes, which were then re-implanted and analysed between 13.5 and 14.5dpc.

The HMG Box is the most conserved region of SRY across species (Whitfield et al., 1993) and where the majority of identified human variants in DSD individuals are located (Supplementary Table 1). As proof of principle of the experimental system, we removed the entire HMG box in the XX Tg line (HMG-Del mutation) (Figure 4). Analysis of gonadal markers via both qPCR and IF showed typical expression of female factor Foxl2 and no obvious testicular phenotype as seen in the non-edited XX Tg embryos.

**Figure 4:**
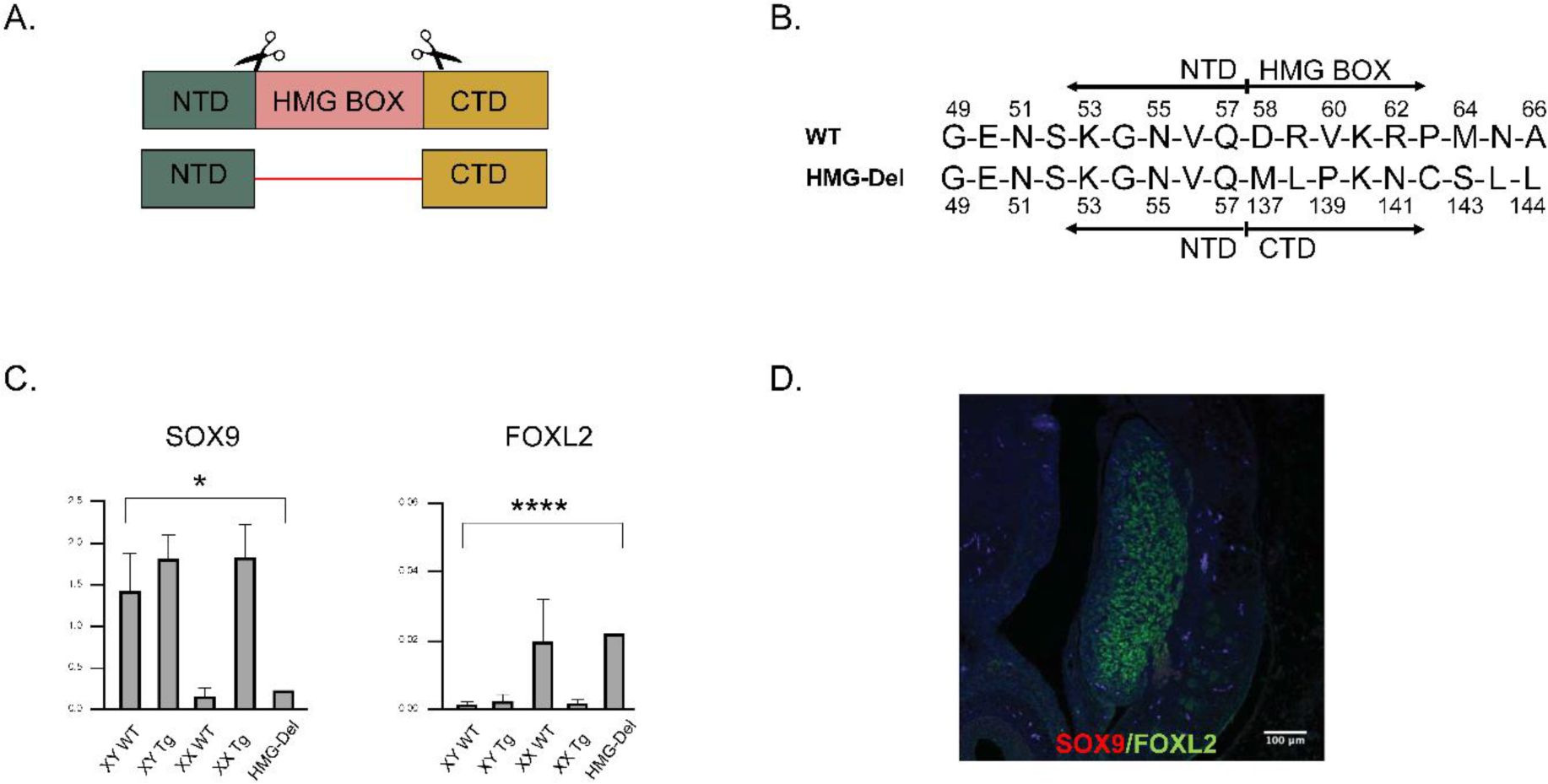
The HMG-Del mutation. **A.** Strategy to generate the HMG-Del variant via CRISPR in XX Tg mice. **B.** Detail of modification in the WT and mutant *SRY*sequences. **C.** *SOX9* and *FOXL2* levels as measured by qPCR of the embryonic gonads. * = p<0.05; **** = p<0.0001 **D.** IHC of the resulting gonad at 14.5 dpc, stained by SOX9 (red) and FOXL2 (green).

### Modification of the HMG box from AA131 to AA136 inactivates *SRY* function and prevents testis development (“Sml-HMG-Del-Mis” mutation)

SRY must first enter the nucleus to regulate downstream gene expression, and the two nuclear localization signals (NLS) within the HMG box are believed to be necessary (Südbeck and Scherer, 1997, Poulat et al., 1995). We designed a guide RNA to create a double strand break to interrupt the RPRRK domain of the NLS at position AA131 (“Sml- HMG-Del-Mis” alteration). A 9 bp deletion, followed by a 6 bp insertion altered this domain, while keeping the open reading frame intact (Figure 5). The resulting gonads appeared phenotypically female, and qRT-PCR confirmed that *Foxl2* was expressed at levels of a typical wild-type XX ovary. *Sox9* mRNA levels were higher in Sml-HMG-Del-Mis XX Tg compared to XX wild type gonads, but not as elevated as wild-type XY controls or non-edited XX Tg testes. However, SOX9 protein was not detected by IF analysis; the gonads instead showed high levels of FOXL2 throughout. These results demonstrated that the elimination of the NLS inactivated the ability of *hSRY* to trigger testis development.

**Figure 5:**
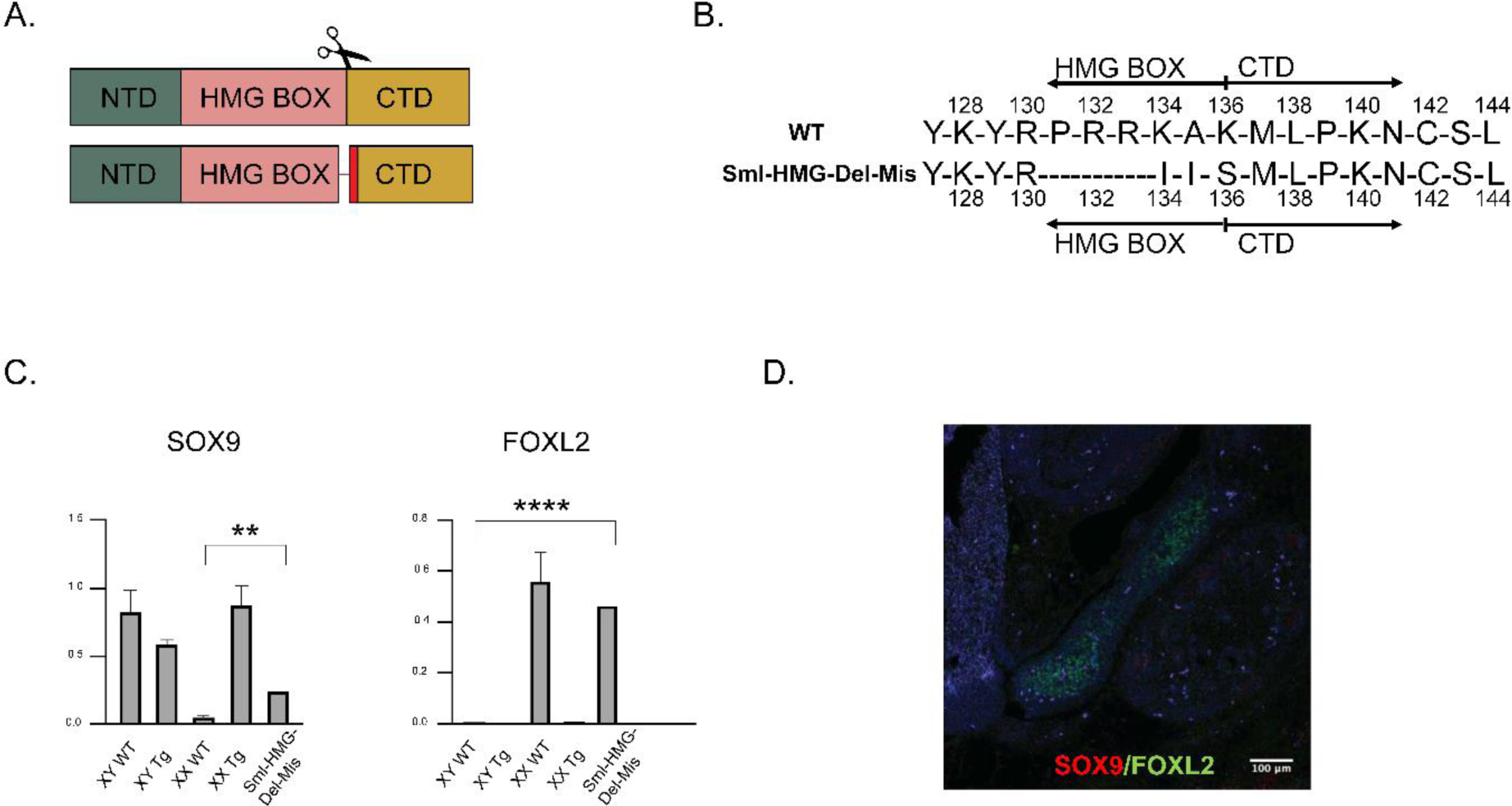
The Sml-HMG-Del-Mis mutation. **A.** Strategy to generate the Sml-HMG-Del-Mis variant via CRISPR in XX Tg mice. **B.** Detail of modification in the WT and mutant *SRY* sequences. **C.** *SOX9* and *FOXL2* levels as measured by qPCR of the embryonic gonads. ** = p<0.01; **** =p<0.0001 **D.** IHC of the resulting gonad at 14.5 dpc, stained by SOX9 (red) and FOXL2 (green).

### Deletion of the C-terminal domain from AA163 to AA165 results in ovotestis development (“C-Del” mutation)

A well-studied pathogenic variant in *hSRY* found in DSD is an early termination at AA163 (Tajima et al., 1994). This variant eliminates the last 41 amino acids of the protein, leaving a CTD that is only 26 amino acid long. We targeted this position using a guide RNA to generate an indel and introduce a new stop codon (C-Del mutation). The resulting variant deleted codons at position 163-165 and altered the codon at position 166 (Figure 6). The resulting gonad appeared similar to a testis during gross examination. However, further characterization revealed an ovotesticular phenotype with high expression of both *Sox9* and *Foxl2 by* qRT-PCR. IF analysis confirmed that SOX9 was confined to the centre, whereas FOXL2 was expressed throughout the gonad, as expected in ovotestes. These findings showed that loss or alteration of the C-terminal domain residues 163-166 reduced but did not eliminate hSRY function.

**Figure 6:**
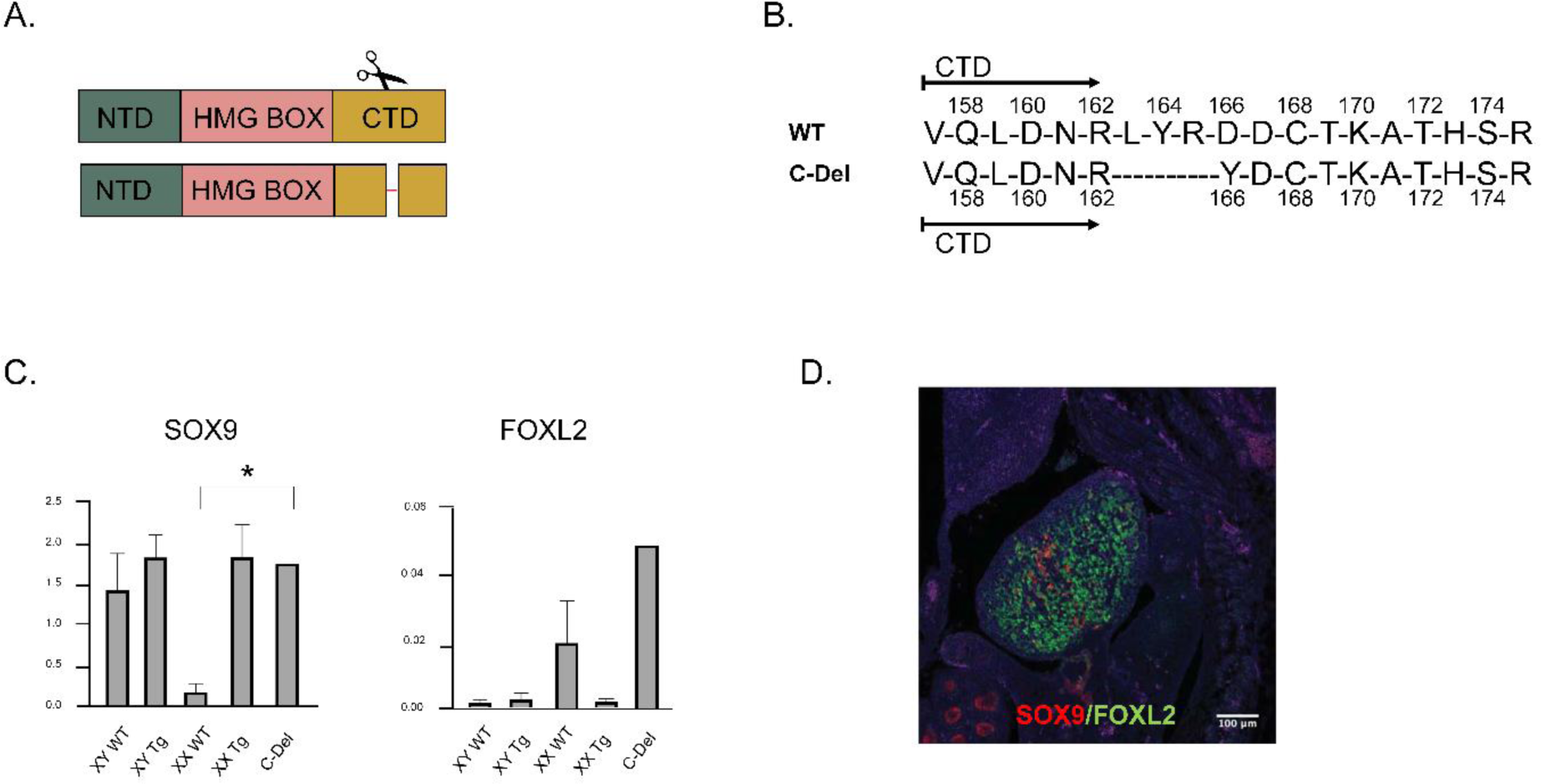
The C-Del mutation. **A.** Strategy to generate the C-Del variant via CRISPR in XX Tg mice. **B.** Detail of modification in the WT and mutant *SRY* sequences. **C.** *SOX9* and *FOXL2* levels as measured by qPCR of the embryonic gonads. * = p<0.05. **D.** IHC of the resulting gonad at 14.5 dpc, stained by SOX9 (red) and FOXL2 (green).

### Early termination of the N-terminal domain inactivates SRY function and prevents testis development (“N-STOP” mutation)

Nine distinct variants of the *SRY* NTD have been reported in DSD, with over half of these resulting in early stop codons (Supplementary Table 1). We generated a change closely resembling the G54Fs*59 variant recently reported (Xue et al., 2019). A 1bp insertion at AA56 led to a frameshift and early termination codon at AA59, the second AA of the HMG Box (N-STOP mutation) (Figure 7). As expected, due to loss of the *hSRY* DNA binding domain, the resulting gonad developed into an ovary. qRT-PCR showed upregulation of *Foxl2* and downregulation of *Sox9* in mutant XX Tg gonads compared to non-edited XX Tg testes. IF staining confirmed FOXL2 expression throughout the ovary and no visible SOX9 protein was detected. These results confirm that changes that remove the HMG box result in complete inactivation of hSRY, leading to complete sex-reversal.

**Figure 7:**
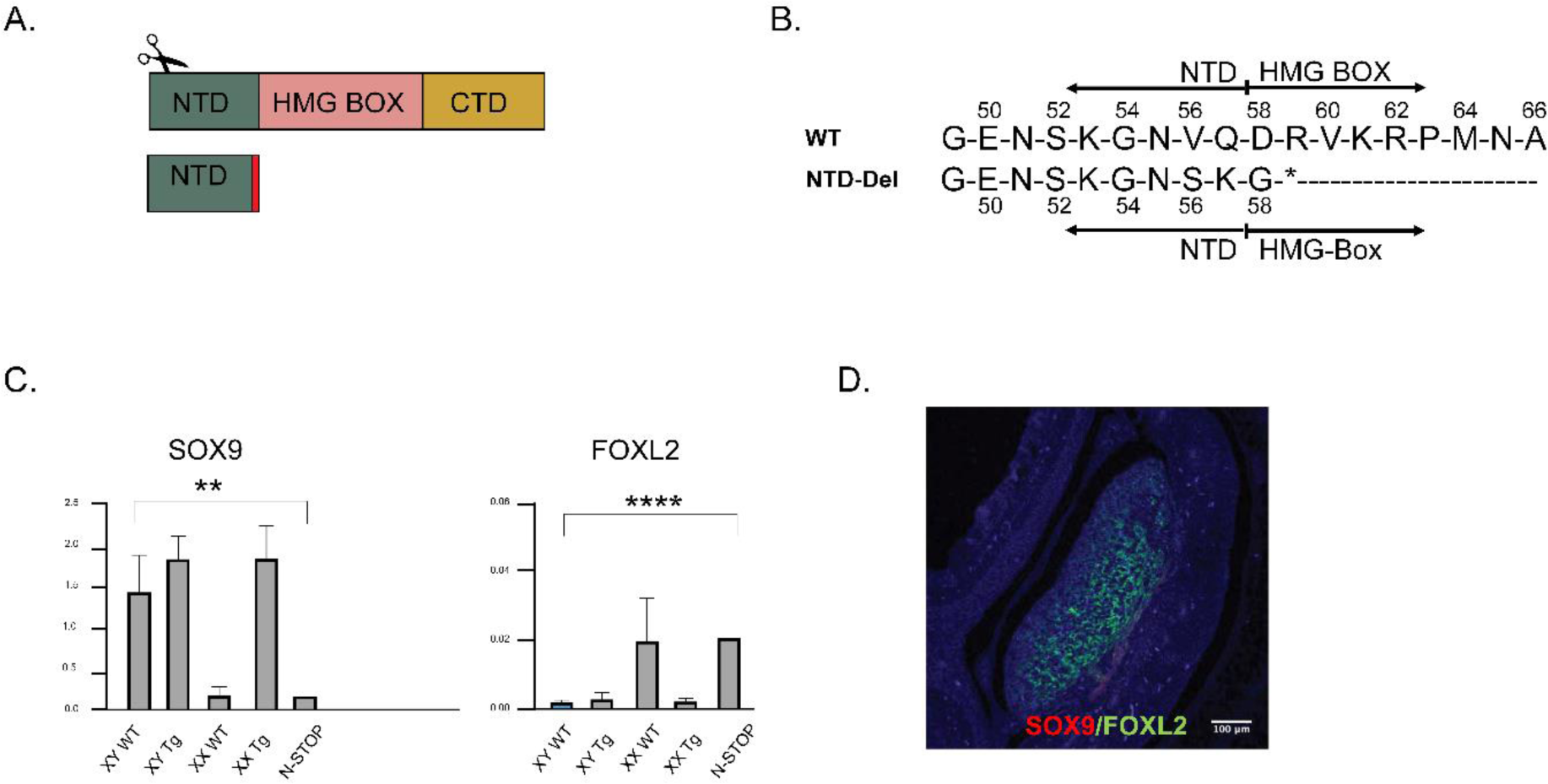
The N-STOP mutation. **A.** Strategy to generate the N-STOP variant via CRISPR in XX Tg mice. **B.** Detail of modification in the WT and mutant *SRY* sequences. **C.** *SOX9* and *FOXL2* levels as measured by qPCR of the embryonic gonads. ** p<0.01; **** p<0.0001 **D.** IHC of the resulting gonad at 14.5 dpc, stained by SOX9 (red) and FOXL2 (green).

### *SRY* variants fail to fully activate SOX9 expression

SRY binds to the *SOX9* enhancers TESCO and Enh13 in mice, activating expression within the bipotential gonad (Gonen et al., 2017, Croft et al., 2018). To first quantify the binding of three of the *hSRY* variant forms generated here (HMG-Del, SmlHMG-Del-Mis, C-Del) *in vitro* DNA occupancy at the *SOX9* enhancers was measured via ChIP-qPCR (Figure 8A, Supplementary Figure 2). As expected, no ChIP-qPCR signal was observed on transient transfection of an empty plasmid, or of an inactive variant of SRY (I68A, a “cantilever” variant that abolishes specific DNA binding (Weiss et al., 1997, Phillips et al., 2004). Each of the variants generated in the transgenic XX mice exhibited reduced TESCO/Enh13 enhancer occupancies (HMG-Del SRY: zero; C-Del mutation: ∼50%; Sml-HMG-Del-Mis mutation: ∼20% relative to wild type (Figure 8A).

**Figure 8:**
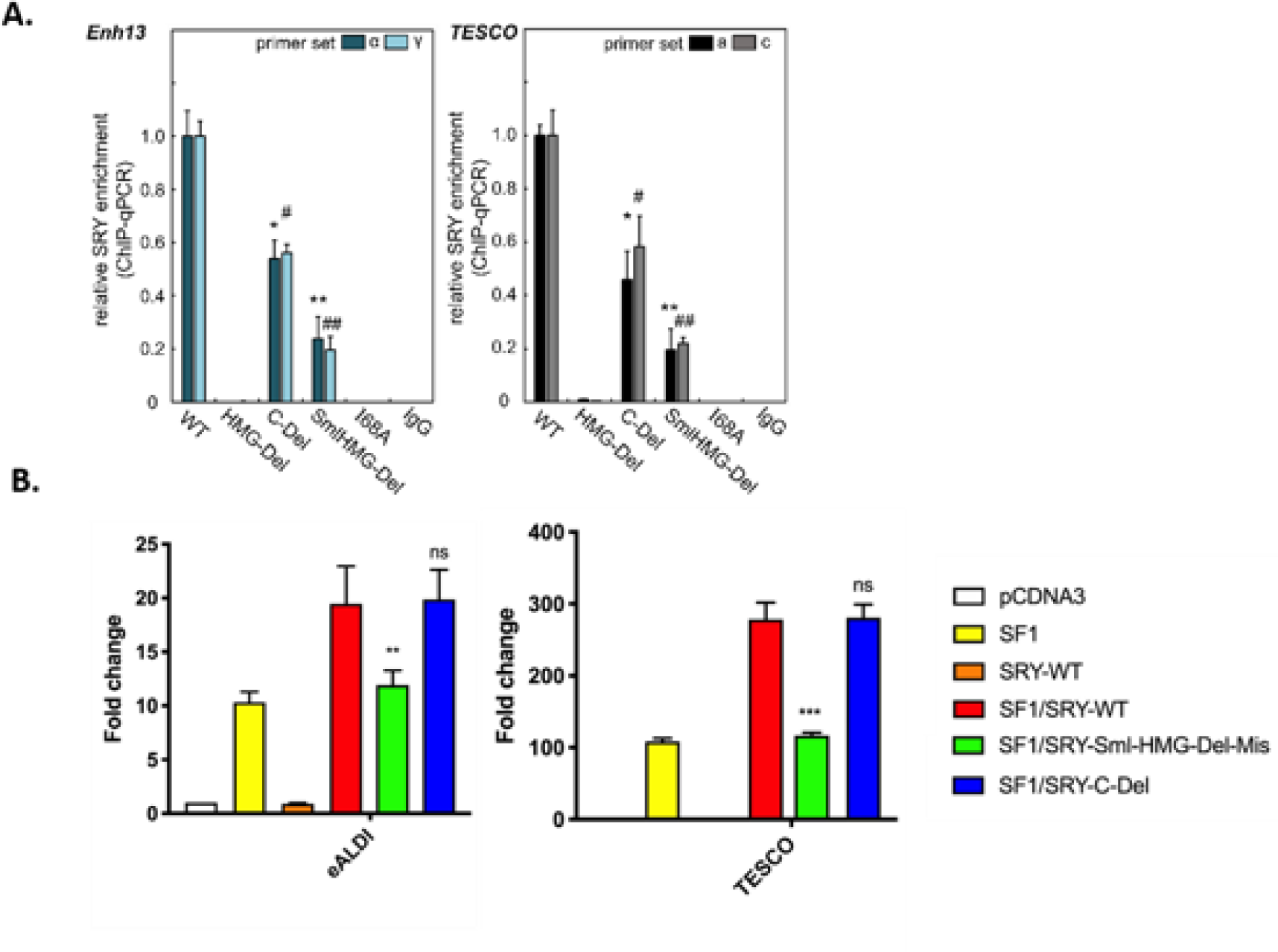
SRY variant activity *in vitro*. **A.** *SOX9* enhancer occupancies associated with various SRY mutations. SRY-*Enh13* (left) and SRY-*TESCO* (right) complexes were probed by ChIP assays and quantified by qPCR; results are summarized by histogram. The experiments was conducted in triplicate on three biological replicates in CH34 cells. Results were normalized relative to WT SRY (far left in each panel). Control lanes I68A and IgG indicate inactive SRY variant and non-specific control. Error bars represent standard deviation. *p < 0.05, primer set α or a, # p < 0.05, primer set γ or c, **p < 0.01, primer set α or a, ## p < 0.01, primer set γ or c. **B.** *In vitro* luciferase assays to assess transcriptional activation of the *SOX9* eALDI and TESCO enhancers in COS7 cells by SF1 and either wild-type or mutant SRY. The data is represented as mean fold change of luciferase activity, relative to that obtained by the empty pCDNA3 vector, from four independent assays, each with two technical replicates. Error bars represent standard error of the mean (SEM). Ratio paired parametric t-tests were performed and differences in luciferase activity compared to SF1 are depicted above corresponding bars. Differences in luciferase activity compared to SF1/SRY-WT are depicted on lines above the bars. *p < 0.05, **p < 0.01, ***p < 0.001, ****p < 0.0001, ns: not significant, WT: wild type.

Next, activation of the TESCO enhancer, as well as the human eALDI enhancer, was assessed via luciferase assays (Croft et al., 2018) (Figure 8B). This was performed in COS7 cells with the addition of SF1 and compared WT SRY with the C-Del and Sml-HMG-Del-Mis variants. As expected, WT SRY alone did not activate either TESCO or eALDI enhancer, but with the addition of SF1 was able to drive luciferase expression fully. The C-Del variant was still able to activate both TESCO and eALDI enhancers to similar levels as WT SRY, while the Sml-HMG-Del-Mis showed ∼30 % activity at TESCO and ∼50% activity at ALDI. We engineered additional *SRY* variants (Supplementary Figure 3). These also resulted in SRY inactivation but were less informative than those presented above.

Thirdly, we assessed the effect of SRY on expression of endogenous *SOX9* gene in an optimized cell culture assay. Two cell lines were exploited for this purpose, CH34 (Haqq and Donahoe, 1998), derived from the rat gonadal ridge, and LNCap (Horoszewicz et al., 1983), derived from human prostate cancer cells. These lines were chosen as they both exhibit SRY-directed transcriptional activation of *SOX9*, unlike most other mammalian cell lines. We (Y-S.C., M.W.) have refined this assay over a number of years to provide a high level of reproducibility and a low level of variability (see Supplementary Figures 4 and 5, and Supplementary Methods for details); a critical feature is low level of transfected gene expression (10^3^-10^4^ SRY molecules per cell). The assay circumvents the need for a reporter construct and relies instead on an active chromatin structure at the endogenous target gene, as well as baseline expression of cofactors presumably required for assembly of an SRY-dependent enhanceosome.

Using this assay, we found that HMG-Del *SRY* was devoid of activity and generated similar *SOX9* mRNA levels to the empty vector (Figure 9A, blue bars). The C-Del variant showed 50% activation of SOX9, and the Sml-HMG-Mis-Del 30% relative to wild type. In each case, reduced activation of *SOX9* was associated with impaired expression of male-pathway-related genes downstream of *SOX9*, including *Sox8*, *Ptgds* and *Fgf9*, whereas expression of house-keeping genes (*Gapdh*, *β-actin*, *Tbp* and *Yy1*) was unaffected (Supplementary Figure 6). Fold-changes in *SOX9* transcription on transient transfection of the *SRY* constructs were similar in the two unrelated *SRY*-responsive cell lines (Figure 9A) despite marked differences in baseline *SOX9* mRNA abundances (Supplemental Figures 4 and 5).

**Figure 9:**
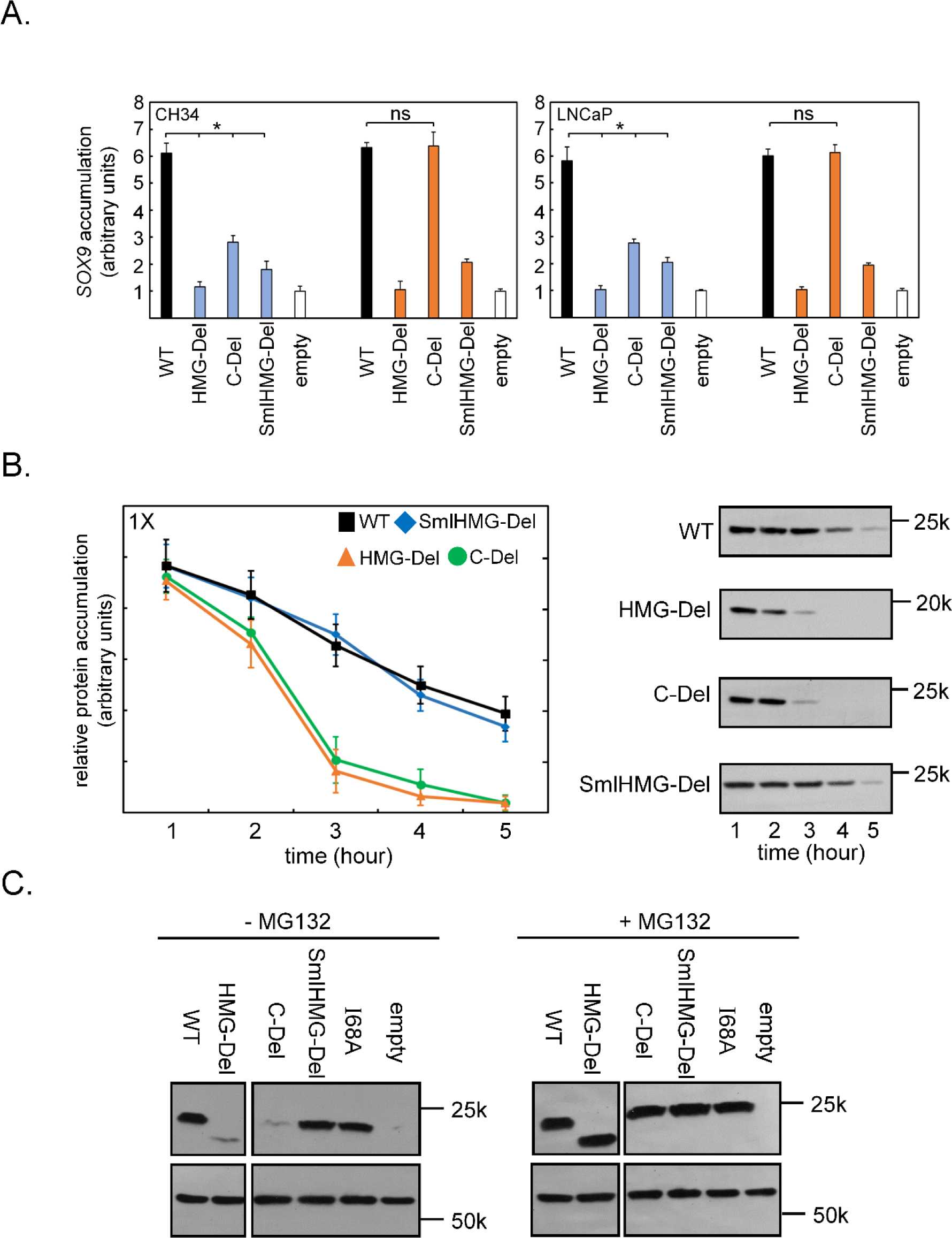
Proteasomal degradation of SRY variants. **A.** *Sox9*expression after transfection of SRY variants. The normalized qPCR data were obtained in the absence (left, blue) or presence (right, orange) of MG132 in CH34 and LNCaP cells. Vertical axis provides fold-changes in abundance of SOX9 mRNA (a ratio); see Supplemental Figures 4 and 5 (in absence/presence of MG132) for individual terms in ratio. The experiment was performed in triplicate on three biological replicates. **B.** SRY protein degradation assays in CH34 cell lines. The semilog plot of SRY concentration as a function of time (left) following cycloheximide arrest of new protein translation. Western blots (WBs; right) exhibited similar time courses. **C.** Addition of proteasome inhibitor MG132 restores SRY protein accumulation in cells to WT levels. WB employed anti-HA antiserum; loading-control α-tubulin is shown in lower box.

### Blocking protein degradation rescues SRY activity in C-terminal but not HMG box or N-terminal alterations

To investigate whether the observed inactivation of *hSRY* was due to accelerated degradation of the altered forms of SRY, new protein translation was blocked using cycloheximide, and perdurance of the epitope-tagged protein was evaluated by western blot (Figure 9B). Both C-Del and HMG-Del showed accelerated degradation, while the Sml- HMG-Del-Mis was comparable to wild type SRY.

When we applied the proteasomal inhibitor MG132 to block degradation, the perdurance of C-Del *hSRY* was fully restored, whereas no effect was observed in HMG-Del and Sml-HMG- Del-Mis *hSRY* (Figure 9C). MG132 also restored the ability of the C-Del to activate *Sox9* expression to wild type levels (Figure 9A). These findings suggest that proteasomal degradation was responsible for the pattern of decreased functional activity of C-Del *hSRY*, in turn suggesting that the CTD of *hSRY* is not only involved in transcriptional activation (Dubin and Ostrer, 1994), but is also required for protein stability.

## Discussion

In the present study, we used CRISPR to modify *hSRY* in a transgenic mouse model and study the relationship between the resulting variants and SRY function. Variants in individuals with DSD have been reported in each of the three domains of SRY: NTD, HMG Box, and CTD. Whereas the majority of these variants are found in the HMG Box, several missense and frameshift changes have been identified within the flanking domains, providing genetic evidence of their importance for full SRY function. The critical role of the HMG domain in DNA binding and bending, and nuclear importation, has been demonstrated both *in vivo* and *in vitro* (Sánchez-Moreno et al., 2008, Kurtz et al., 2021). As proof of principle, we successfully removed the HMG box while keeping the transcript in frame. The HMG box-deleted hSRY failed to bind to the sex-specific mouse *Sox9* enhancers and activate *Sox9* transcription, resulting in ovarian development in XX Tg mice. These experiments also confirmed that CRISPR HDR repair is achievable through electroporation without using any microinjection techniques, allowing a simplified workflow.

The SRY HMG box contains two nuclear localization signals (designated nNLS and cNLS) that are required for nuclear import of SRY (Poulat et al., 1995, Südbeck and Scherer, 1997, Forwood et al., 2001). The nNLS binds calmodulin (Sim et al., 2005) and Exportin4 (acting in nuclear import (Sim et al., 2005), whereas the cNLS binds importin-B. We mutated cNLS by replacing essential amino acids with missense variants. This resulted in inactivation of hSRY function and rescue of ovarian development. A different change causing a premature stop codon at AA132 also interrupted the cNLS and resulted in the same phenotype (Supplementary Figure 3). We expected that deletion of only one NLS would partially impair nuclear entry but retain specific DNA-binding/bending activity. Indeed, the Sml-HMG-Del-Mis hSRY was found in both cell lines to exhibit ∼20% activity (as evaluated by *Sox9* enhancer occupancy and *Sox9* transcriptional activation; Figures 8 and 9), and this reduction in functionality could not be rescued by proteasome inhibition. We propose that this was due to the reduced ability of the protein to enter the nucleus, rather than its lack of DNA-binding activity. Consistent with this hypothesis, *in vitro* analyses of a variant SRY HMG box containing a clinical mutation in the cNLS (R133W) showed DNA-binding affinity equivalent to wild type (Harley et al., 2003, Li et al., 2001). The *in vitro* inactivity of HMG-Del *hSRY* and reduced activity of Sml-HMG-Del-Mis *hSRY* at equivalent levels of protein expression (Figure 9) are likely due to qualitative effects, i.e., loss of specific DNA binding in the former and impaired nuclear entry in the latter. Our results support the critical role of the cNLS for the correct SRY nuclear shuttling and function during gonadal development.

The functional role of SRY’s CTD has not been fully understood. In DSD individuals, six variants have been reported in the CTD of *SRY* to date (Shahid et al., 2004, Sánchez-Moreno et al., 2009, Baldazzi et al., 2003, Shahid et al., 2005, Tajima et al., 1994, Özen et al., 2017). Affected individuals usually display female external genitalia and streak gonads. The CTD spans from AA137 to AA204 and was shown to possess transcriptional activation activity (Dubin and Ostrer, 1994). The most distal variant associated with DSD was found at AA178 and caused an early stop codon. The individual with this variation, who identified as male, exhibited ambiguous genitalia, a single testicle, and a contralateral streak gonad (Özen et al., 2017). This report suggests that while detrimental, the mutation does not completely abolish SRY function. Rare variants C-terminal to AA178 can be found in the Gnomad database (Karczewski et al., 2020). However, these variants are not associated with impaired gonadal development, suggesting that they do not significantly affect SRY activity. These reports seem to be consistent with the CTD’s transcriptional activity being located within the first ∼40 amino acids of this domain.

Interruption of the C-terminal AA163-166 sequence in our mouse model resulted in the development of ovotestes, which contained both Sertoli and granulosa cells. Ovotesticular DSD is the rarest form of DSD with an incidence of less than 1/20000 (Scarpa et al., 2017). Currently, only two cases of ovotestis have been reported in non-mosaic 46,XY individuals, and both mutations occurred at position AA60 in the HMG box (Berta et al., 1990, Hiort et al., 1995). *In vitro* analyses of these mutations showed partial activation of *Sox9,* indicating that SRY maintained some DNA binding activity (Phillips et al., 2011). Similarly, our C-Del *hSRY* retained 50% activity compared to wild type SRY, and its function was fully recovered by proteasome inhibition. Therefore, we hypothesize that the sequence around AA163-166 is important for protein stability. The C-Del mutation could also affect the partner binding activity of SRY. SRY is thought to bind the KRAB-O protein, thus recruiting KAP1 to repress transcription of pro-ovarian genes (Oh et al., 2005, Peng et al., 2009). This interaction has been mapped to the region around AA139 to AA162 of SRY, one amino acid before our C- Del alteration (Oh et al., 2005). It is possible that this variant may disrupt the interaction between SRY and KRAB-O, consistent with the inability to repress expression of the ovarian gene *Foxl2*. Our results demonstrate that truncation or internal deletion of the CTD are associated with accelerated protein turnover. However, the C-Del mutation did not disrupt the transcriptional activation capacity of the CTD, which is unlikely to localize within positions 163-166. How the CTD protects SRY from proteasomal degradation is not well understood. It would be of future interest to investigate the critical boundary between stable and unstable CTD deletions, and whether the stabilizing effect of the native CTD is intrinsic to its length or instead mediated by specific binding to a partner protein. As previously mentioned, the mouse *Sry* lacks an homologous CTD but instead contains a large glutamine-rich repeat, which also protects the protein from proteasomal degradation (Zhao et al., 2014a). Analogous protection of human and mouse *Sry* by divergent CTDs likely reflects distinct molecular mechanisms Similar to the CTD, variants within the NTD of SRY are less common than those within the HMG box. Most N-terminal variants lead to early termination upstream of the HMG box, and are therefore uninformative regarding NTD function. Using CRISPR, we introduced an early termination codon at AA59, predictably showing loss of SRY’s ability to direct male sex determination and a lack of *Sox9* activation. Further work will be required to generate more specific and informative NTD variants for study.

## Conclusions

CRISPR technology has dramatically improved our ability to study human disorders using animal models. We have described a new mouse model system that combines zygote electroporation, CRISPR genome-editing and specific *in vitro* assays. In the future, this model can be used to investigate the molecular roles of SRY in initiating testicular differentiation, including studies of DSD variants and general structure-function relationships.

## Methods

### Generation of hSRY transgenic mice

The Wt-hSRY transgenic mouse was generated using the methods outlined in(Zhao et al., 2014b). Briefly, the 3FLAG-hSRY-GFP was cloned into pENTR1-A, and a LR Clonase reaction was performed to introduce the 3FLAG-hSRY-GFP into the PB-Wt1-Dest vector containing the gateway enhanced piggyBAC vector with Wt1 regulatory regions. All animal procedures were approved by the University of Queensland Animal Ethics Committee.

The PB-Wt1-3F-hSRY-GFP and hyPBase mRNA were diluted in water for embryo transfer, 2ng/uL and 2.5ng/uL respectively and micro injected into the pronucleus of B6CBAF1/J zygotes. After microinjection, zygotes grown to the 2-cell stage overnight and transferred into pseudo pregnant foster CD1 mice. From 3 microinjection sessions 60 founder pups were born, of which 5 contained the transgene (3XX and 2XY). Breeding between the founder XY tg and wild type females confirmed germline transmission, and the line was back crossed onto a C57BL6 background.

### Splinkerette PCR

After genomic DNA extraction, splinkerette PCR was performed following the protocol outlined in (Li et al., 2010) Sanger sequencing of products was performed and aligned to regions of the mouse genome using BLAT (Kent, 2002).

### Genotyping

Standard genotyping was performed by DNA extraction with QuickExtract buffer (Lucigen), after ear clipping (pups) or tail clipping (embryo). Sextyping and Tg PCR primers were used to genotype each sample. Sanger sequencing was performed on samples after genotyping with Phusion Polymerase (NEB) and PCR Purification Kit (QIAGEN). Primer sequences listed in Supplementary Table 2.

### Postnatal Reproductive Organ Analysis

Mice of each genotype (XX WT, XX Tg, XY WT, XX Tg) were culled at 6 weeks of age and their reproductive organs dissected for analysis. Ovaries and testes with epididymides were paraffin embedded and stained using the protocols outlined below.

### Haemotoxylin and Eosin Staining

Paraffin embedded postnatal reproductive organs were sectioned at 7uM (Leica HistoCore Multicut Microtome). A Leica HistoCore Pearl tissue processor was used for staining, using standard haemotoxylin and eosin protocols. Slides were mounted with MountQuick (Daido Sangyo) and imaged on an Olympus BX51 Microscope with Olympus DP70 CCD camera.

### CRISPR Electroporation

To generate zygotes for electroporation, 3-week-old C57BL6 females were superovulated with subcutaneous injection of PMSG (5.8 IU) and HCG(7.5IU) 46 hours apart. After HCG injection, females were mated overnight with XY Tg males, and checked for presence of a copulation plug the next morning. Females were culled and oviducts collected and placed in pre-warmed M2 for zygote collection with the addition of hyaluronidase to remove cumulus cells. Zygotes washed 3x in pre-warmed and CO2 equilibrated KSOM and kept at 37°C, 5% CO2.

CRISPR guide RNA and tracRNA must first be ligated to form the duplex cr:tracRNA. The crRNA (IDT) and tracRNA (IDT) were diluted to 200uM in Duplex Buffer (IDT) and mixed at an equimolar concentration to 95°C for 5 min and allowed to cool to RT. This was added to Opti-MEM to a final concentration of 6 μM, and 1 μL of 61 μM Cas9 nuclease (IDT) is added for a final concentration of 1.2 μM. If a HDR template was to be added, this was diluted to 200uM and a final concentration of 20 μM. CRISPR mix was incubated at room temperature for 15-20 min prior to electroporation.

Electroporation was performed using the NEPA21 Electroporator System (NEPAGENE). 5uL of CRISPR mix was added to the chamber of the electrode, and up to 30 zygotes washed in pre-warmed Opti-MEM were added to the mix. The impedance was measured, and liquid was added or removed to ensure this was between 0.18 and 0.21 ohms. The poring pulse (40V, 3.5 ms, 50 ms interval, 4x), and the transfer pulse (5V, 50 ms, 50 ms interval, 5x) was administered to the zygotes, and removed immediately after. Zygotes washed 3 times in KSOM, before overnight incubation at 37°C, 5% CO2. Embryo transfer into pseudo pregnant CD1 foster mothers was performed for 2-cell embryos the next day. Transient embryo retrieval was then performed at 13.5- or 14.5 days gestation.

### Immunostaining

Embryos were fixed overnight in 4% PFA, washed with PBS, dehydrated in EtOH series and embedded in paraffin for sectioning. Sections were taken at 7uM (Leica HistoCore Multicut). For fluorescent immunostaining, slides were dewaxed in xylene and rehydrated with EtOH series (100%, 90%, 80%, 70%, 50%, H_2_O) using an autostainer (Leica HistoCore Autostainer XL). Antigen retrieval was performed with high pH Tris-based Antigen unmasking solution (Vector Labs), then washed (H_2_O, PBTX) and blocked with blocking solution for 4 hours before primary antibody was added and left overnight at 4°C. Primary antibody washed off with PBTX washed, reblocked with blocking solution for 1 hour, and secondary antibody kept on for 4 hours in the dark. Slides washed with PBTX and PBS and mounted with Prolong Gold with DAPI. Confocal imaging performed with Zeiss LSM 710 Microscope and edited with FIJI software.

Primary antibodies used - Rabbit anti-SOX9 (AB5535, Millipore, 1/800), Goat anti-FOXL2 (NovusBio NOVNB1001277, 1/100), Rabbit anti-MVH (ab13840, Abcam, 1/1000), Mouse anti-FLAG (Sigma F3165). Secondary antibodies used - Donkey Anti-Rabbit-568 (a10042, Invitrogen), Donkey Anti-Goat-488 (A11055, Invitrogen), Goat Anti-Mouse-568 (A11003, Invitrogen).

### qRT-PCR

During embryo dissection, one gonad with mesenephros was removed and kept in RNAlater (Qiagen) at -20°C until RNA extraction. RNA extraction performed using RNEasy Micro Kit (QIAGEN), and RNA converted to cDNA with High Capacity RNA to cDNA Kit (Applied Biosystems). Triplicate SYBR qRT-PCR assays were performed on the QuantStudio 7 machine (Applied Biosystems), and relative level of expression was calculated after normalisation to *Tbp*. Primers used are listed in Supplemental Table 2. Statistical analyses performed in Prism.

### Mammalian expression plasmids

Plasmids expressing full-length SRY and variants were constructed by PCR and verified by DNA sequencing. Following the initiator Met, the cloning site encoded a hemagglutinin (HA) tag in triplicate to enable Western blotting (WB) and chromatin immunoprecipitation (ChIP).

### Cell culture

Rodent CH34 cells (Haqq and Donahoe, 1998, Chen et al., 2013) were cultured in Dulbecco’s modified Eagle’s medium (DMEM) containing 5% fetal bovine serum (FBS) at 37°C in 5% CO2. LNCaP cells (ATCC® CRL-1740™) were obtained from ATCC and cultured in DMEM medium containing 10% FBS in 5% CO2.

### Transient transfections

Transfections were performed using Lipofectamine 3000 as described by the vendor (Invitrogen). After 8 hours in Opti-MEM medium, cells were recovered using fresh culture medium with FBS: Dulbecco’s Modified Eagle Medium (DMEM) for CH34 cells and Roswell Park Memorial Institute (RPMI) medium for LNCaP cells. Transfection efficiencies were determined by ratio of green-fluorescent protein (GFP) positive cells to untransfected cells following co-transfection with pCMX-SRY and pCMX-GFP in equal amounts (Chen et al., 2013). Subcellular localization was visualized by immunostaining 24-h post transfection following treatment with 0.01% trypsin (Invitrogen) and plating on 12-mm cover slips. To assess transfection efficiency, cells were visualized via fluorescence microscopy for qualitative assessment of protein expression, morphology, and viability. SRY expression was monitored by WB via its triplicate HA tag.

### Chromatin immunoprecipitation

Cells were transfected with epitope-tagged WT or variant SRY. SRY-expressing cells were cross-linked in wells by formaldehyde, collected, and lysed after quenching the cross-linking reaction. Lysates were sonicated to generate fragments and immunoprecipitated with anti-HA antiserum (Sigma) containing a Protein A slurry (Cell Signaling). A non-specific antiserum (control IgG; Santa Cruz) served as non-specific control. PCR and qPCR protocols were as described (Racca et al., 2016). Quantification of immunoprecipitated enhancer was performed by qPCR (Racca et al., 2016) and representative gel images were collected by Gel Doc (Bio-Rad).

### Cycloheximide assay and western blotting

24-h post transient transfection, cells were split evenly into 6-well plates and treated with cycloheximide to a final concentration of 20 μg/ml in DMEM for the indicated times; cells were then lysed by RIPA buffer (Cell Signaling Technology). Protein concentrations were measured by BSA assay (Thermo); cell lysates were subjected to 4-20% SDS-PAGE and WB using anti-HA antiserum (Sigma-Aldrich) at a dilution ratio of 1:5000; α-tubulin antiserum provided a loading control. For phosphorylation analysis, HA-tagged SRY variants were immunoprecipitated with rabbit polyclonal anti-phosphoserine antiserum (Abcam). WB following 4-20% sodium dodecyl sulfate-polyacrylamide gel electrophoresis (SDS-PAGE) employed HRP-conjugated anti-HA antibody (Roche). Quantification was performed by Image J software.

### Transcriptional activation assay

In the transient transfections the expression plasmid encoding HA-tagged WT SRY or an HA-tagged variant SRY was diluted 1:50 with the parent empty plasmid to reduce protein expression to the physiological range (*ca*. 10^3^-10^4^ protein molecules/cell (Chen et al., 2013)); similar expression levels were verified by anti-HA Western blot. Following transient transfection, cellular RNA was extracted and converted to cDNA by using vendor’s protocol (Bio-Rad). SRY-mediated transcriptional activation of *SOX9/Sox9* was measured in triplicate by qPCR as described (Chen et al., 2013). Primer sequences for all of the tested genes were applied as described (Chen et al., 2013, Racca et al., 2016). *Tbp*, encoding the specific 5’-TATAADNA-binding subunit of TFIID, was used as an internal control; experiments included three technical replicates of each of three biological replicates. For further detail see Supplementary Methods.

### Luciferase Assays

To assess mutant or wild-type SRY activity, *in vitro* luciferase assays were performed as described in Croft et al. (2018). Briefly, COS7 cells were seeded in a 96-well plate for 2-4 hours in DMEM with 10% foetal bovine serum at 37°C with 5% CO2. At 80–90% confluency, 75 ng of the enhancer-luciferase reporter constructs pGL4-eALDI, pGL4-TESCO or empty pGL4.10 (Promega;(Croft et al., 2018)) were individually transfected into the cells in each well in duplicates, together with 100 ng each of the following transcription factors in pCDNA3.1-based expression constructs: SF-1, wild-type/mutant SRY or SF-1 + wild-type/mutant SRY. The pRL-TK plasmid (Promega; 15 ng) expressing Renilla luciferase was included as an internal control. The cells were transfected using Lipofectamine 2000 (Thermo Fisher Scientific) for 24 hours. Cells were then lysed and firefly and Renilla luciferase activities assayed using the Dual Luciferase Reporter Assay System (Promega).

Four independent assays were performed, and firefly luciferase activity was normalised against that of Renilla luciferase. Fold change was calculated by normalising luciferase activity against the average of the empty vector pCDNA3.1. Data was represented as mean fold change of the four biological replicates with standard error of the mean (SEM). Statistical significance was assessed by performing two-tailed Student’s paired t-tests using Graphpad Prism V7.

## Acknowledgements

We thank Johnny Huang for performing microinjections and the UQ Transgenic Animal Service of Queensland (TASQ) for performing embryo transfer. We thank P.K. Donahoe (Massachusetts General Hospital, Boston, MA) for kindly providing the CH34 cell line. This work was supported by research grants from the National Health and Medical Research Council of Australia, and the Distinguished Professor endowment at Indiana University (MAW).

## Abbreviations

ALDI: Alternate Long-Distance Initiator
ChIP: chromatin immunoprecipitation
CTD: C-terminal domain
DSD: differences (or disorders) in sexual differentiation
FBS: fetal blood serum
GFP: green fluorescent protein
HA: haemagglutinin
HMG: high mobility group
HRP: horse radish peroxidase
hSRY: human SRY
mSRY: mouse Sry
NTD: N- terminal domain
PCR: polymerase chain reaction
qRT-PCR: quantitative real-time reverse- transcriptase PCR
SDS-PAGE: sodium dodecyl sulfate-polyacrylamide gel electrophoresis
*SRY/Sry*: sex-determining region of the Y chromosome
TESCO: testis-specific enhancer core element
TF: transcription factor
Tg: transgenic
WB: western blot. Amino acids are designated by standard one- or three-letter codes. Names of genes (or acronyms) are italicized.

