## Supplemental Methods for "Generation and mutational analysis of a transgenic mouse model of human SRY"

**Supplementary Methods**

Plasmids

The backbone of the SRY expression plasmid is a pCMX, containing an CMV promotor and encoding an N-terminal 3X-HA epitope tag following the initiator Met. Plasmids expressing full-length SRY and variants were constructed by PCR and verified by DNA sequencing.

Transient Transfections

To obtain the best of transfection efficiency, we use the recommended protocol provided by Thermo-Fisher Scientific:

https://www.thermofisher.com/us/en/home/references/protocols/cell-culture/transfection-protocol/lncap-cells-protocol.html.

Transfection protocols were performed with a mixture of the pCMX empty parent (0.98 µg) and pCMX plasmids with target SRY-encoded genes (0.02 µg), with the overall transfected mass of plasmids retained at 1 µg as standardly recommendeded for 1 million cells. We designate this dilution protocol "50X plasmid dilution conditions." Such dilution of the expression plasmid attenuates cellular expression of protein by 320-fold (maximum feasible dilution) to provide a physiologic intracellular TF concentration (details was published by Lodish, H., et al. (2000) Molecular Cell Biology 4th Ed; our protocol was developed as described in Chen, Y-S., et al. PNAS (2013)). This protocol mitigates potential artifacts due to TF overexpression without affecting the transfection efficiency in either cell line. Our purpose is to obtain transfected cell expressing appropriate protein copies (1000-10000 protein molecules per cell; $10^3$ – $10^4$) without reducing the relative percentage of transfected cells.

Transcriptional Activation Assay

Following transient transfection, SRY-mediated transcriptional activation of the endogenous SOX9 or Sox9 gene was measured by qPCR.

Sample preparation followed the Bio-Rad protocol entitled “Two-step RT-qPCR.” In this two-step method, RNA is first transcribed into cDNA in a reaction using reverse transcriptase. An aliquot of the resulting cDNA can then be used as a template for multiple qPCR reactions. (In the one-step method, RT and qPCR are performed in the same tube.)
In two-step RT-qPCR, the RT step can be primed either with oligo(dT) primers, random primers, a mixture of the two, or gene-specific primers. One-step RT-qPCR must be performed using gene-specific primers and can be achieved either by using Thermus thermophilus (Tth) polymerase, a DNA polymerase with inherent RT activity, or by a two-enzyme system combining a reverse transcriptase with a thermostable DNA polymerase.

qPCR Plate Protocol Optimizing Consistency and Reproducibility

When comparing qPCR results from different source materials, a reference gene that is expressed at the same level in all samples enables normalization with respect to differences in overall starting levels of RNA or small sample-to-sample differences in reaction efficiency. Researchers typically pick a common reference gene, such as GAPDH or TBP, based on their stable expression (ideally at the same level in all samples and under all conditions). Although reference gene expression can sometimes vary (for example, GAPDH expression in liver cells is modulated by hormonal signaling), we have carefully analyzed GAPDH or TBP as “house-keeping genes” in the CH34 and LNCaP cell lines using the geNorm method (Hellemans et al. (2007)).

In the present qPCR assays there were two sources of potential variability to be assessed: (1) biological variability and (2) technical variability. Biological variability is due to inherent differences between individual organisms, tissues or cell culture samples. Technical variability is introduced through experimental factors such as variability in volume of liquid pipetted (pipetting error) or variability in the filling or sampling individual wells in a plate. The present protocol employed three biological replicates and three technical replicates per biological replicate. Thus, in the technical replicates three wells for the same aliquot was used to account for technical variability; any measured differences can be attributed to technical (or experimental) variability. To obtain the three biological replicates per assay, we used multiple 6-well plates with different sets of cells treated with same set of SRY variants. A helpful factor in our experimental design was designing a simple plate well-assignment scheme as follows.

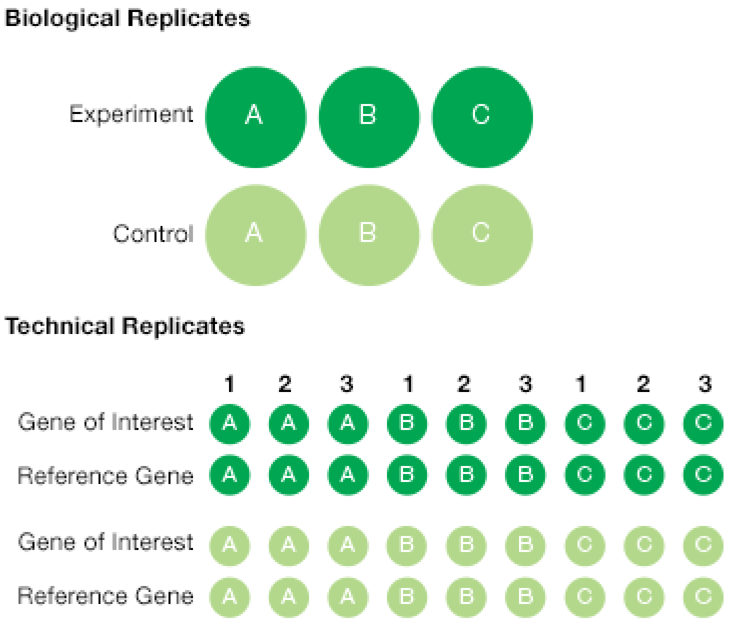


The above schematic figure represents experimental replicates of our standardized plate plan in accordance with the recommendations of the vendor (Bio-Rad).

In additional, we routinely ran a common sample with a known quantity of DNA on each plate as a bridge for consistency among multiple plates. This control sample enabled detection and normalization for run-to-run variability.

<u>Organization of Data Sets</u>

To prepare the figure in the main text (Figure 9A), the data were represented fold as fold-differences. Fold-values pertain to the ratio of endogenous target gene expression (SOX9 or Sox9) on transient transfection of an SRY construct to its baseline expression on transient transfection of an empty vector (1 μg parent plasmid per transfection reaction). Because the fold-difference is a ratio X/Y, there can be (and usually are) cell-line-specific differences in individual X and Y values even when the ratio is similar between cell lines. For example,

CH34 cells and LnCaP cells exhibit different transfection efficiencies as well as species-specific and lineage-specific differences in baseline gene expression and chromosomal architecture. Fortuitously, their respective endogenous Sox9 and SOX9 genes exhibit similar patterns of gene-specific histone marks, in each case associated with amenability to SRY-driven transcriptional activation and in each case associated with similar patterns of homologous sex-specific enhancer occupancies.
