## Supplemental Figures for "Generation and mutational analysis of a transgenic mouse model of human SRY"

**Supplementary Figure 1: Localization of the hSRY Transgene**

Location of the piggyBAC vector (YourSeq) as determined by Splinkerette PCR is on chromosome 8 (qA1.1), within the intergenic region upstream of SOX1.

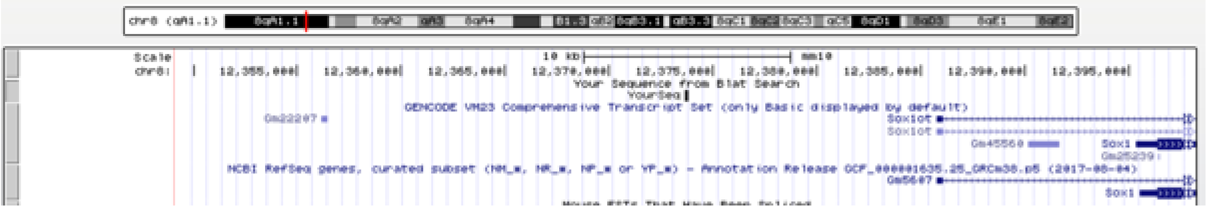

**Supplementary Figure 2: Sox9 enhancers**

*Sox9* gene is regulated by a far upstream enhancer element (*Enh13;* aqua) and *TES* core elements (*TESCO*; light grey). Potential SRY binding sites are present in *Enh13* fragments 1 and 3 (related to primer sets α and β for ChIP assay) and *TESCO* fragments 4 and 8 (primer sets a and c for ChIP assay).

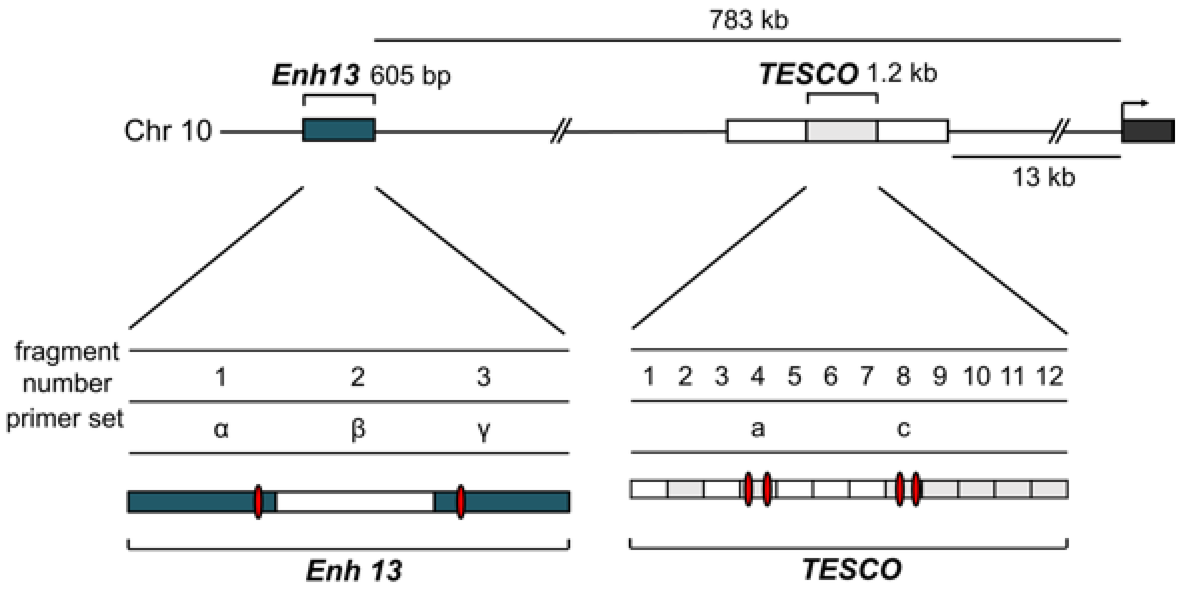

**Supplementary Figure 3: Additional variants generated by CRISPR.**

**A.** Variant #1 contained a deletion within the NTD abolishing the ATG site. The closest methionine, and presumably new start site was at position 64. Variant #2 contained an early termination within the HMG box at position 132. Variant #3 featured 4 missense amino acids in positions 56-59, followed by an early stop codon. Variant #4 contained a deletion of 66 amino adics at position 134-200. Variant #5 contained a deletion within the CTD at position 193-197, followed by 24 missense amino acids. Immunohistochemistry for SOX9 (red) and FOXL2 (green) showed extensive staining of ovarian marker FOXL2 throughout the entire gonads. Scale bar, 100 μm. Sequence variations are shown compared to WT (bold-top) **B.** qRT-PCR for SOX9 and FOXL2 was performed at E14.5 for variations #1,#3,#4,and #5. Error bars represent the standard error of the mean (SEM). **C.** qRT-PCR for SOX9 and FOXL2 was performed at E13.5 for variation #2. **D.** *In vitro* luciferase assays to assess transcriptional activation of the *SOX9* eALDI (orthologous to mouse Enh13) and TESCO enhancers in COS7 cells by SF1 and either wild-type or *CRISPR* variations of SRY. The data are represented as mean fold change of luciferase activity, relative to the empty pCDNA3 vector, from four independent assays, each with two technical replicates. Error bars represent standard error of the mean (SEM). Ratio paired parametric t-tests were performed and differences in luciferase activity was compared to SF1/SRY-WT.

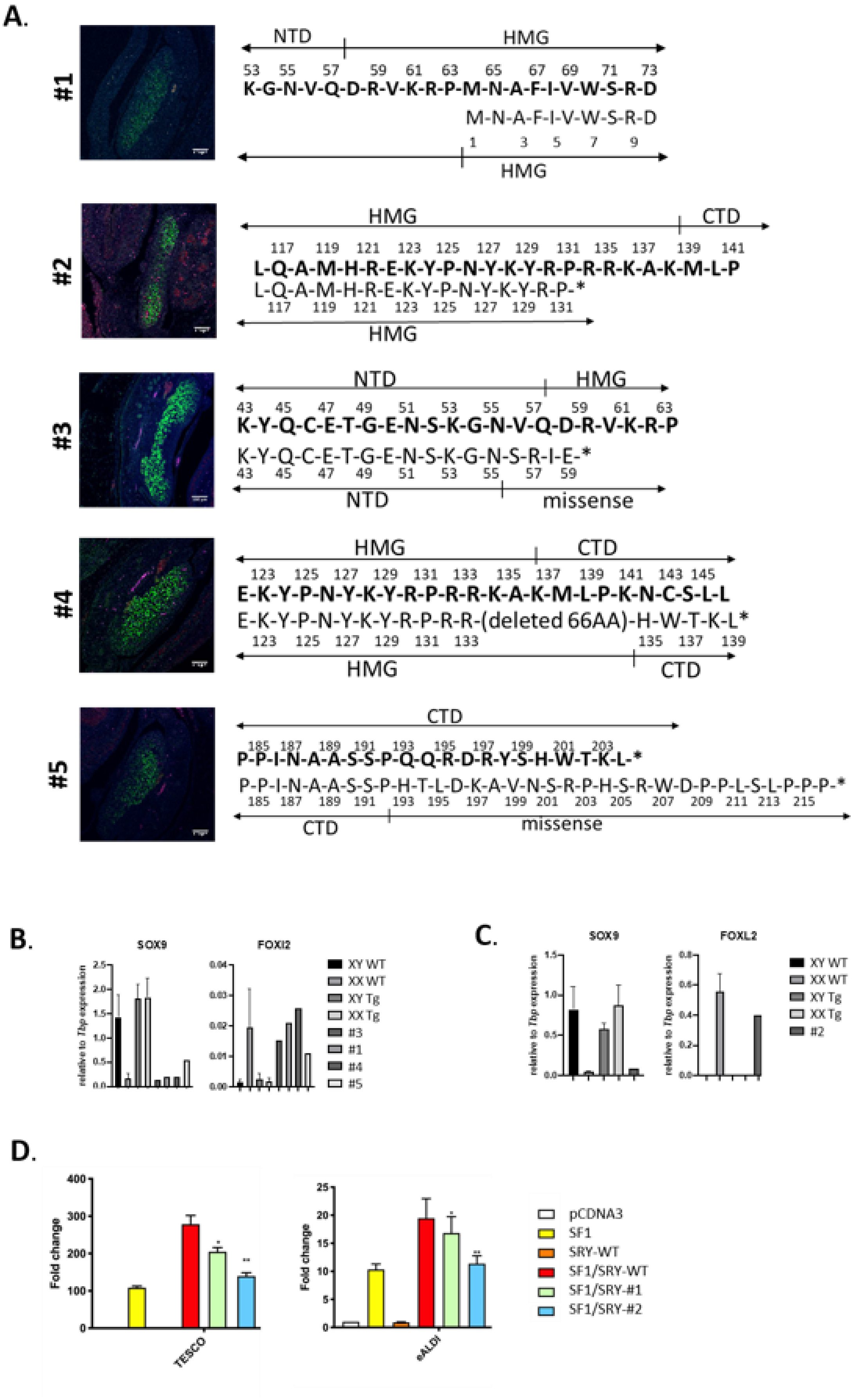
A.
#1
NTD
HMG
53 55 57 59 61 63 65 67 69 71 73
K-G-N-V-Q-D-R-V-K-R-P-M-N-A-F-I-V-W-S-R-D
M-N-A-F-I-V-W-S-R-D
1 3 5 7 9
HMG
#2
HMG
CTD
117 119 121 123 125 127 129 131 135 137 139 141
L-Q-A-M-H-R-E-K-Y-P-N-Y-K-Y-R-P-R-R-K-A-K-M-L-P
L-Q-A-M-H-R-E-K-Y-P-N-Y-K-Y-R-P-*
117 119 121 123 125 127 129 131
HMG
#3
NTD
HMG
43 45 47 49 51 53 55 57 59 61 63
K-Y-Q-C-E-T-G-E-N-S-K-G-N-V-Q-D-R-V-K-R-P
K-Y-Q-C-E-T-G-E-N-S-K-G-N-S-R-I-E-*
43 45 47 49 51 53 55 57 59
NTD
missense
#4
HMG
CTD
123 125 127 129 131 133 135 137 139 141 143 145
E-K-Y-P-N-Y-K-Y-R-P-R-R-K-A-K-M-L-P-K-N-C-S-L-L
E-K-Y-P-N-Y-K-Y-R-P-R-R-(deleted 66AA)-H-W-T-K-L*
123 125 127 129 131 133 135 137 139
HMG
CTD
#5
CTD
185 187 189 191 193 195 197 199 201 203
P-P-I-N-A-A-S-S-P-Q-Q-R-D-R-Y-S-H-W-T-K-L-*
P-P-I-N-A-A-S-S-P-H-T-L-D-K-A-V-N-S-R-P-H-S-R-W-D-P-P-L-S-L-P-P-P-*
185 187 189 191 193 195 197 199 201 203 205 207 209 211 213 215
CTD
missense
B.
SOX9
FOXl2
relative to Tbp expression
XY WT
XX WT
XY Tg
XX Tg
#3
#1
#4
#5
C.
SOX9
FOXL2
relative to Tbp expression
XY WT
XX WT
XY Tg
XX Tg
#2
D.
Fold change
TESCO
eALDI
pCDNA3
SF1
SRY-WT
SF1/SRY-WT
SF1/SRY-#1
SF1/SRY-#2

**Supplementary Figure 4: Transfection Assay Methodology**

Expression of wild-type (WT) SRY or variant SRY constructs in two SRY-responsive mammalian cell lines, one derived from the rat XY gonadal ridge (CH34; panel A) and the other from a human XY metastatic prostate-cancer deposit (LNCaP; panel B). Histograms provide respective Sox9/SOX9 mRNA abundances (as measured by qPCR) following normalization with respect to the abundance of a housekeeping mRNA (GAPDH) in each cell line. Data were obtained in the absence of proteasomal inhibitor MG132. Arrows at bottom highlight inequivalent scales between cell lines, ascribed to cell-line-specific differences in transient-transfection efficiency, species of origin (rat vs human), developmental lineage and stage (embryonic gonadal ridge vs adult prostate) and mechanism of immortalization (Ras-transformed vs metastic malignancy). Despite such differences, fold-differences are similar as shown in Figure 9A in main text.  SRY constructs (left to right in each panel): wild-type (WT), HMG-Delta (HMG-Δ), C-Delta (C-Δ) and Sml-HMG-Delta (Sml-HMG-Δ) as defined in main text.

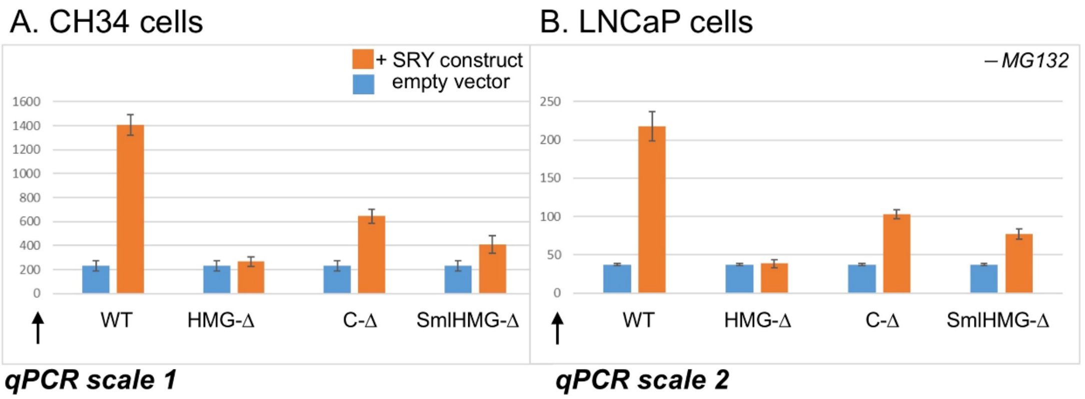

**Supplemetary Figure 5: Protein Degradation Assay Methodology**

Addition of proteasomal inhibitor MG132 differentially affects SRY-directed transcriptional activation of respective endogenous Sox9/SOX9 genes depending on SRY modification. Relative to patterns of gene expression in absence of MG132, proteasomal inhibition significantly enhances the activity only of SRY variant C-Delta (C-Δ), suggesting that the C-terminal non-box segment of human SRY augments in cellular lifetime. A mammalian XY cell line derived from the rat XY gonadal ridge (CH34; panel A) and a human XY cell line derived from a metastatic prostate-cancer deposit (LNCaP; panel B) give rise to cell-line-specific scales of mRNA abundances (arrows at bottom) but similar fold-changes in Sox9/SOX9 gene expression as shown in Figure 9A in main text. SRY constructs (left to right in each panel): wild-type (WT), HMG-Delta (HMG-Δ), C-Delta (C-Δ) and Sml-HMG-Delta (Sml-HMG-Δ) as defined in main text.

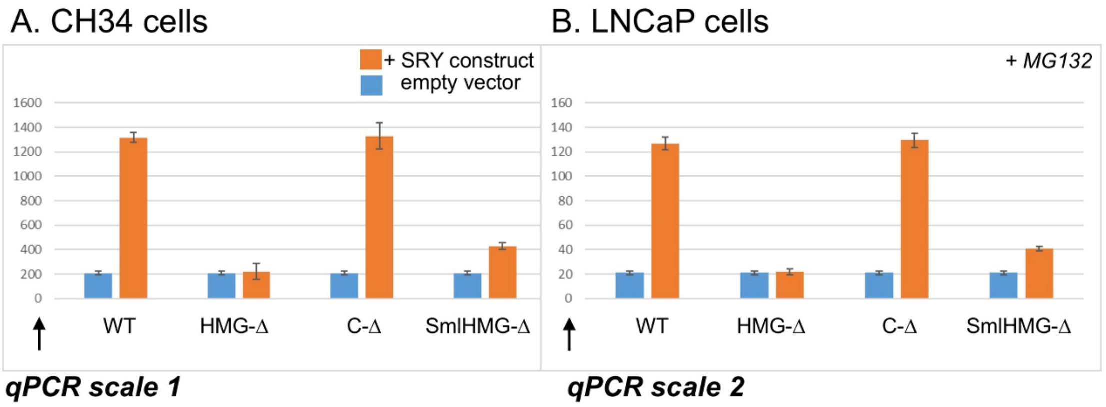

**Supplementary Figure 6: Downstream responses to SRY variants**

Survey of mRNA expression of male-differentiation regulation genes (*Sox*8, *Sox9*, *Ptgds*, and *Fgf9*), and endogenous *rat Sry*. Controls were provided by unrelated housekeeping genes. *Inset box*, statistical significance at the level $p<0.05$ (Wilcox test) for SRY-dependent transcriptional activation (*). Experiments were performed in triplicate on each of three biological replicates.

**Supplementary Table 1: SRY pathogenic variations identified in human individuals with DSD (Excel spreadsheet)**

Data taken from literature of DSD genomic analysis where a pathogenic variation was identified within SRY. The colour of the codon number denotes the location (green=NTD, pink=HMG Box, yellow=CTD). Where information is not available within the report, it is left blank within the table.

**Supplementary Table 2 (Excel spreadsheet)**

Oligonucleotide sequences used in experiments.
